# SMURF1/2 are novel regulators of WNK1 stability

**DOI:** 10.1101/2024.07.31.606092

**Authors:** Ankita B. Jaykumar, Sakina Plumber, Derk Binns, Chonlarat Wichaidit, Katherine Luby-Phelps, Melanie H. Cobb

## Abstract

Angiogenesis is essential for remodeling and repairing existing vessels, and this process requires signaling pathways including those controlled by transforming growth factor beta (TGF-β). We have previously reported crosstalk between TGF-β and the protein kinase With No lysine (K) 1 (WNK1). Homozygous disruption of the gene encoding WNK1 results in lethality in mice near embryonic day E12 due to impaired angiogenesis and this defect can be rescued by endothelial-specific expression of an activated form of the WNK1 substrate kinase Oxidative Stress-Responsive 1 (OSR1). However, molecular processes regulated via a collaboration between TGF-β and WNK1/OSR1 are not well understood. Here we show that WNK1 interacts with the E3 ubiquitin ligases SMURF1/2. In addition, we discovered that WNK1 regulates SMURF1/2 protein stability and vice versa. We also demonstrate that WNK1 activity regulates TGF-β receptor levels, in turn, controlling TGF-β signaling.

## Introduction

Transforming growth factor beta (TGF-β) signaling modulates vascular remodeling [1]. Homozygous disruption of the gene encoding the protein kinase With No lysine (K) 1 (WNK1) results in defective angiogenesis around embryonic day E12 [2,3]. WNK1 is the most ubiquitously expressed among a family of four related atypical protein-serine/threonine kinases, known for their unique catalytic lysine site [4]. WNK1 binds and catalyzes the phosphorylation of substrates, Oxidative Stress-Responsive 1 (OSR1) and STE20/SPS-1-related proline-alanine-rich kinase (SPAK, *STK39*) [5,6,7,8,9]. We previously reported crosstalk among WNK1 and multiple molecules involved in regulating TGF-β signaling pathways [10,11].

SMURF (SMAD Ubiquitination Regulatory Factor) 1/2 are WW-domain-containing enzymes belonging to the NEDD4 (Neural precursor cell Expressed, Developmentally Down-regulated 4) family of E3 ubiquitin ligases. SMURFs mediate endothelial-mesenchymal transition during cardiovascular development [18,19,21,22,23,24,25,26,27]. SMURF1 controls cellular responsiveness to the TGF-β/SMAD2 pathway [27]. TGF-β initiates redistribution of the TGF-β receptor type II into tight junctions which leads to recruitment of SMURF 1 to cause disintegration of tight junctions, a process critical to regulate cell migration and angiogenesis [12,13]. SMURF2, on the other hand, downregulates TGF-β signaling by targeting itself, SMAD2/3 transcription factors and the TGF-β receptor I, also known as, Activin receptor-like kinase 5 (ALK5) for degradation [18,19, 21, 22, 23, 24, 25 26, 27]. This process is critical to regulate TGF-β signaling-dependent processes such as cell migration and angiogenesis [18-27]. Previous work from our laboratory showed that WNK1 is involved in TGF-β pathway-dependent modulation of SMAD2/3 protein stability [10,11]. However, mechanisms underlying how WNK1 modulates TGF-β signaling and processes that, in turn, regulate WNK1 stability during TGF-β signaling are unknown.

Occludin is a tight junction protein which is important for regulating tight junction integrity [14,15,16]. Furthermore, recent studies show that occludin is also involved in endothelial vascular remodeling and angiogenesis [17]. We previously found that occludin interacts with OSR1 to enable tight junction turnover in a WNK1-dependent manner [10]. Our findings suggested intimate connections between TGF-β pathway molecules with the WNK1/OSR1 pathway to control angiogenesis. These findings lead us to speculate that WNK1 may modulate several TGF-β-dependent functions and that regulation of WNK1 protein stability is an important determinant of TGF-β signaling output.

WNK1 protein turnover is regulated by several E3 ligases. WNK1 undergoes ubiquitination followed by degradation through the proteasome upon recruitment to cullin3 (CUL3)-containing E3 ligase complex [28]. Another study from our group reported that ubiquitin-protein ligase E3 component n-recognin 5 (UBR5) interacts with WNK1 and its deficiency results in increased WNK1 protein [29]. A previous study showed that WNK1 isoforms are substrates of NEDD4-2, another member of the NEDD4 family of E3 ubiquitin ligase [30]. WNK1 binds to and phosphorylates NEDD4-2 [20,31,32]. NEDD4-2 negatively regulates TGF-β signaling by ubiquitin-mediated degradation of SMAD2 and TGF-β type I receptor [33]. Therefore, regulation of NEDD4-2 by WNK1 may also be involved to precisely control TGF-β signaling output to modulate cellular processes underlying angiogenesis. This led us to speculate that there may be other E3 ligases involved in the TGF-β signaling pathway that regulate WNK1 protein stability.

In this study, we identify molecular events that underlie WNK1-dependent inputs to specificity of the TGF-β pathway in endothelial cells. We show interactions between WNK1 and SMURF1/2 and report that WNK1 forms discrete signaling microdomains (sometimes referred to as WNK bodies [20]) for reciprocal regulation of WNK1 and E3 ubiquitin ligase SMURF1/2 turnover. In addition, we discover complex inter-regulation between WNK1 and SMURFs during TGF-β signaling. These findings add to the conclusion that WNK1 collaborates with downstream TGF-β signaling components SMURF1/2 to regulate and fine-tune processes involved in TGF-β signaling such as turnover of TGF-β receptors in a context-dependent manner.

## Results

### WNK1 colocalizes with SMURF1/2

Given the involvement of WNK1 in TGF-β-SMAD signaling [10,11], we examined actions of WNK1 on TGF-β-pathway-dependent functions of SMURF1/2 in endothelial cells. First, we tested whether either SMURF1 or 2 co-localize with WNK1 in primary human umbilical vein endothelial cells (HUVECs). Interestingly, we found that a fraction of WNK1 and SMURF1 as well as SMURF2 were observed in a small number of large punctate structures distinct from their distributions elsewhere in cells [26] (**Figure 1A, 1B**).

**Figure 1:**
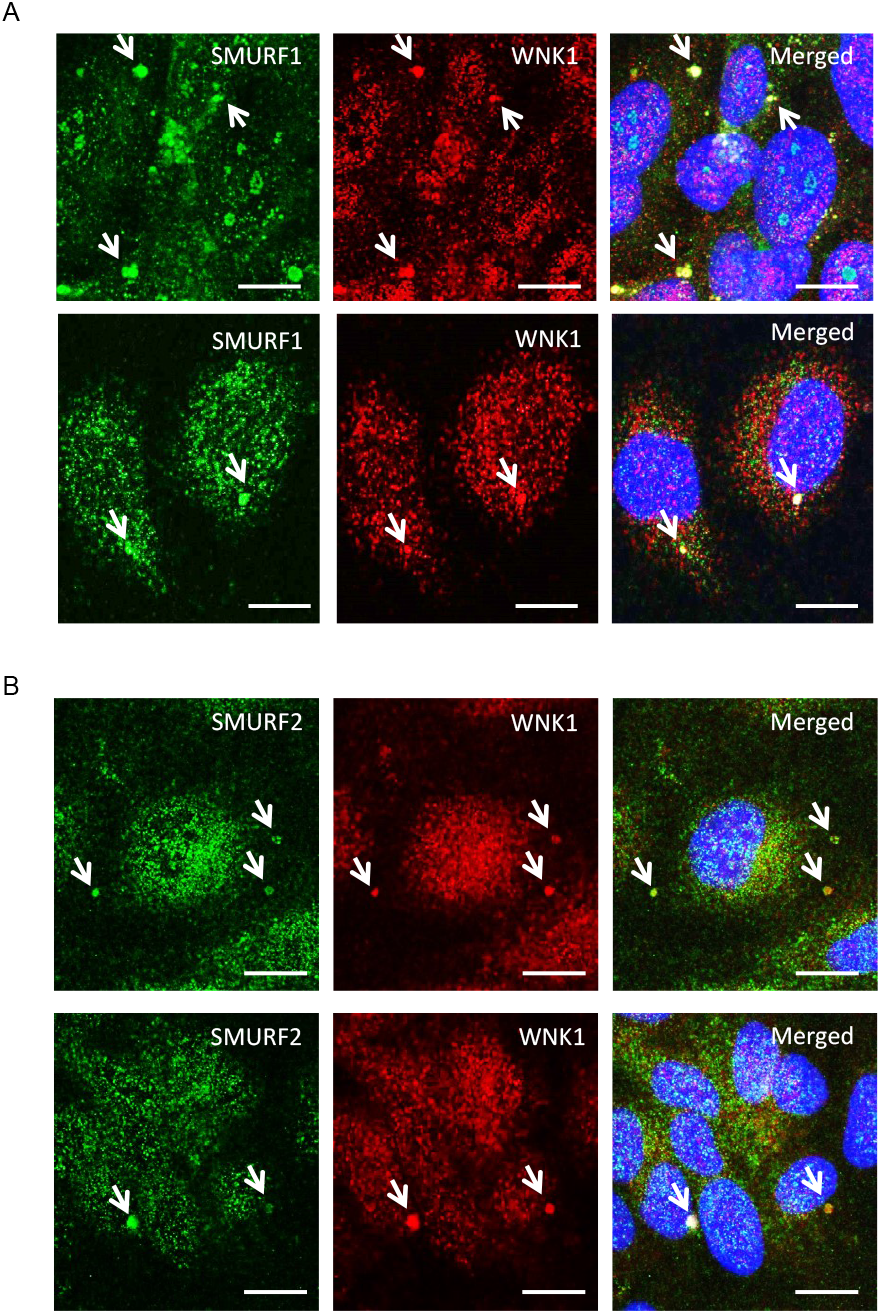
WNK1 colocalizes with SMURF1/2. **A and B**) Representative confocal images of immuno-fluorescently labeled endogenous WNK1 (red), SMURF1/2 (green) and nucleus (DAPI: blue) in primary HUVECs upon 1-2 h TGF-β (10 ng/ml) stimulation. Merged panel (yellow) shows co-localization between WNK1 and SMURF1/2 in large punctate structures. Scale bar = 20 µm; n=3.

### WNK1 kinase activity regulates association of WNK1/OSR1 with SMURF2

Given the colocalization between WNK1 and SMURF1/2, we asked whether WNK1 interacts stably with SMURF1/2 in HUVECs and found SMURF2 in endogenous immunoprecipitates of WNK1 (**Figure 2A**). We found that the WNK1-regulated kinase OSR1 also weakly co-immunoprecipitated with SMURF2 (**Figure 2B**). OSR1 contains a conserved C-terminal (CCT) domain, which can bind substrates and other proteins via short conserved RFxV motifs [34,35]. A blocking peptide can interfere with OSR1 CCT interactions with RFxV motif-containing proteins such as occludin [10]. We then asked whether the interaction with SMURFs is mediated via the OSR1 CCT domain and SMURF RFxV motifs. We found marginal decreases in OSR1 co-immunoprecipitating with SMURF2 in primary HUVECs upon co-incubation with the blocking peptide (**Figure 2B**). We previously found that WNK1/OSR1 regulate the turnover of tight junctions [10]. Therefore, we asked whether SMURF2 interacts with the tight junction protein occludin and found that the interaction between occludin and SMURF2 was enhanced upon inhibition of human dermal microvascular endothelial cells (HDMECs) with the pan-WNK inhibitor WNK463 [10]. This interaction was also diminished upon co-incubation with the blocking peptide (**Figure 2C**). These observations suggest that OSR1 interacts with occludin and weakly with SMURF2 via the CCT domain and that WNK activity regulates the interaction between WNK1/OSR1 and SMURF2.

**Figure 2:**
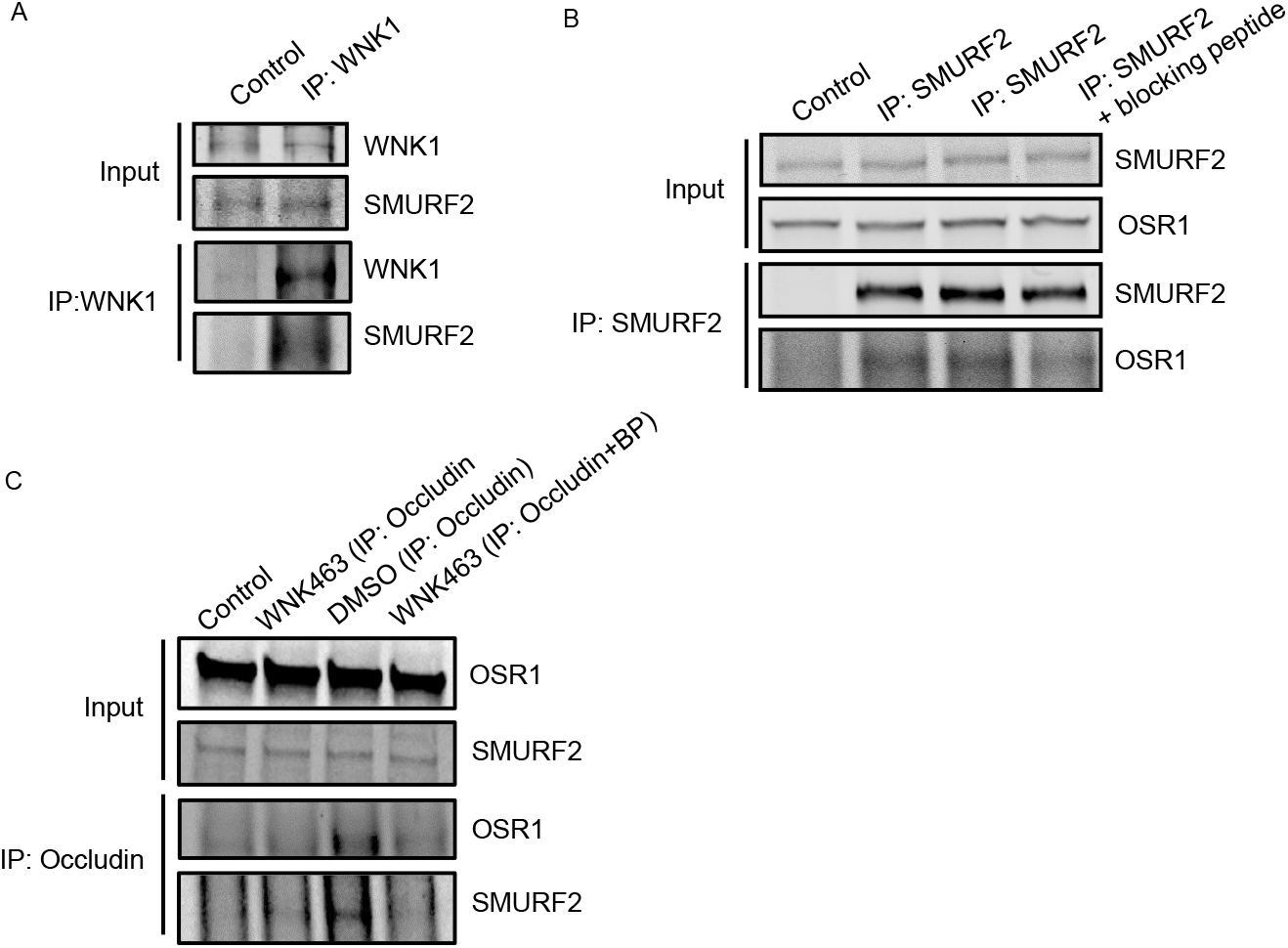
WNK1 kinase activity regulates association of WNK1/OSR1 with SMURF2. **A**) Representative Western blot show endogenous co-immunoprecipitation of SMURF2 with WNK1 show interaction between SMURF2 and WNK1 in HUVECs; n=3. **B**) Representative Western blot show endogenous co-immunoprecipitation of OSR1 and SMURF2 in HUVECs which is diminished upon co-incubation with the blocking peptide SAGRRFIVSPVPE (100 µM); n=3. **C**) Representative Western blot show endogenous co-immunoprecipitation of OSR1 and SMURF2 with occludin in HUVECs upon treatment with WNK463 (1 µM) which is diminished upon co-incubation with the blocking peptide SAGRRFIVSPVPE (100 µM); n=3.

### WNK1/OSR1 regulates SMURF1/2 and vice versa

Given the interaction between WNK1 and SMURFs, we then asked if WNK1 impacts amount of SMURFs. We found that depletion of WNK1 decreased steady state SMURF2 protein expression (**Figure 3A, 3B, 3D, 3E**). In addition, depletion of WNK1 prevented the expected decrease in steady-state SMURF1 protein. However, upon co-treatment with the proteasomal inhibitor MG132 (10 µM), we observed a decrease in SMURF1 protein expression (**Figure 3A, 3C**). Treatment of cells with the SMURF1 inhibitor A01 (2 µM) [36] lead to a significant increase in WNK1 expression (**Figure 3D, 3F**). The amount of SMURF2 protein expressed in the absence of added ligands or inhibitors was related to WNK1 expression (**Figure 3D, 3F, 3G**). In contrast, depletion of OSR1 caused no differences in amounts of either SMURF1 or SMURF2 (**Figure 3A, 3B, 3C, 3G, 3H**). Decreased SMURF2 protein expression was observed only in OSR1-depleted cells that were also treated with either the proteasomal inhibitor MG132 or the SMURF1 inhibitor A01 (**Figure 3A, 3B, 3C, 3G, 3H**). In contrast, SMURF1 protein expression increased in OSR1-depleted cells that were also treated with MG132 (**Figure 3A, 3C**). These results suggest a complex inter-regulation among WNK1, OSR1 and SMURF1/2.

**Figure 3:**
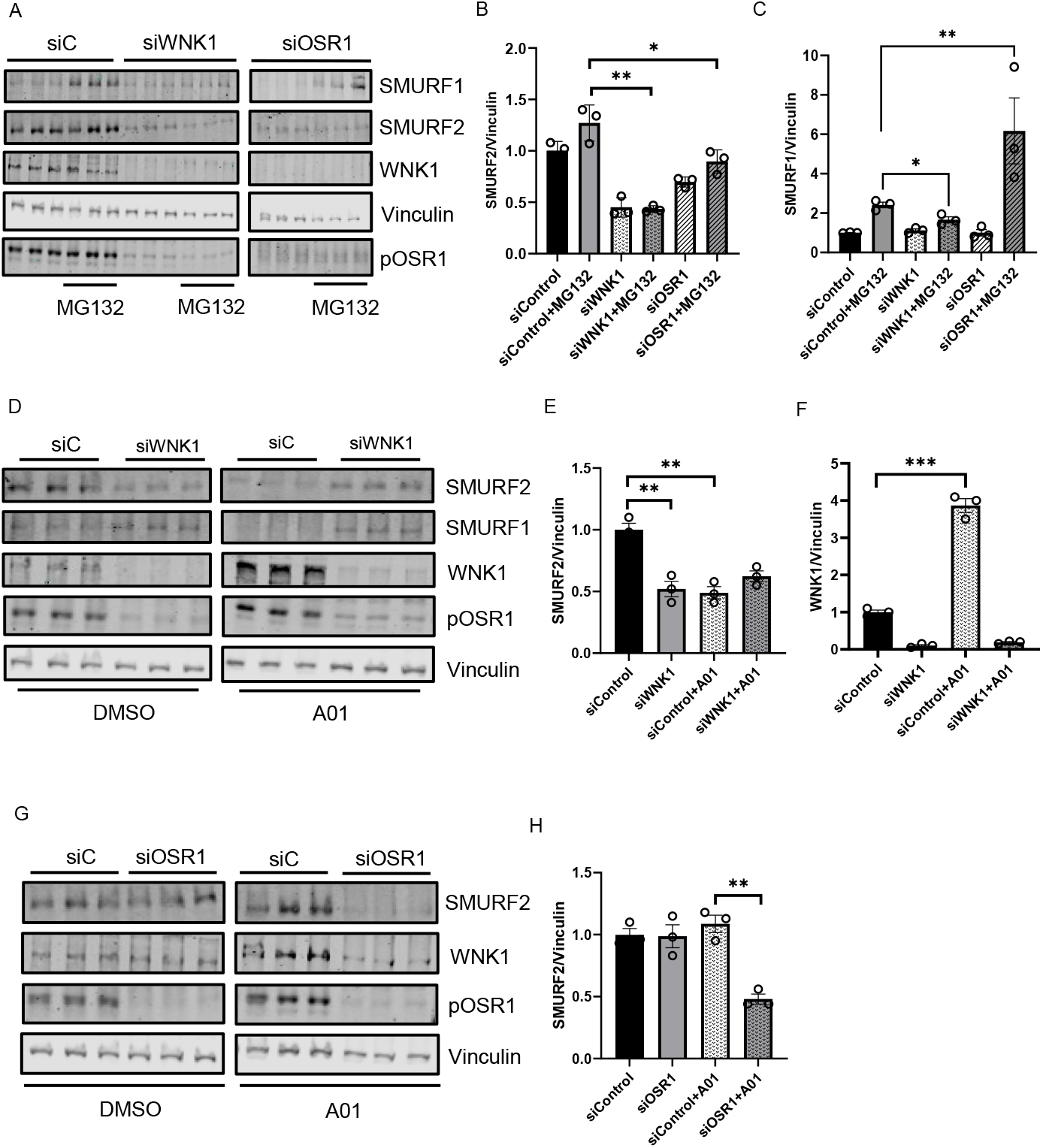
WNK1/OSR1 regulates SMURF1/2 and vice versa. **A**) Representative Western blot show SMURF1/2 expression upon WNK1 or OSR1 depletion by siWNK1 or siOSR1, respectively in HDMECs. It shows increase in SMURF1 levels upon treatment with proteasomal inhibitor MG132 (10µM) for 6 hours which is prevented upon WNK1 or OSR1 depletion. **B**) Corresponding quantification of ‘A’ show decreased SMURF2 levels upon WNK1 and OSR1 depletion compared to siControl; n=3. **C**) Corresponding quantification of ‘A’ show decreased SMURF1 levels upon WNK1 and OSR1 depletion compared to siControl; n=3. **D**) Representative Western blot show SMURF1/2 expression upon WNK1 depletion in HDMECs co-treated with SMURF1 inhibitor A01 (2 µM). **E**) Corresponding quantification of ‘D’ show decreased SMURF2 levels upon WNK1 depletion similar to treatment with the SMURF1 inhibitor A01 (2µM) alone compared to DMSO or siControl; n=3. **F**) Corresponding quantification of ‘D’ show increased WNK1 upon treatment with SMURF1 inhibitor A01 (2µM) alone compared to DMSO or siControl; n=3. **G**) Representative Western blot show SMURF1/2 levels upon siRNA depletion of OSR1 in HDMECs. **H**) Corresponding quantification of ‘G’ show decreased SMURF2 levels with siOSR1 treatment followed by SMURF1 inhibitor A01 (2µM) treatment overnight compared to siControl or DMSO control; n=3. Data are represented as Mean±SE; analyzed by unpaired two-tailed Student’s t-test or one-way ANOVA. *p<0.05, **p<0.005.

### WNK1 kinase activity mediates SMURF2-dependent regulation of ALK5

We asked whether the kinase activity of WNK1 is important for regulation of SMURF2 and found that treatment with WNK463 decreased SMURF2 protein expression, similar to the effect of WNK1 knockdown (**Figure 4A**). We also found that baseline SMURF2 protein expression was enhanced by treatment with the proteasomal inhibitor MG132; WNK463 co-treatment efficiently decreased SMURF2 protein expression (**Figure 4B, 4C**). These results suggest that the observed effects on SMURFs are dependent on WNK1 activity. Previously, we found that WNK463 decreased expression of the type 1 TGF-β receptor ALK1 [10]. Degradation of the TGF-β type I receptor, ALK5, is facilitated via recruitment of the SMURF2 complex [12,13,18,19]. Given the regulation of WNK1 by SMURF2 and effects of WNK inhibition of ALK1, we asked whether WNK1 kinase activity also regulates ALK5. We found that treatment with WNK463 did, in fact, decrease ALK5 protein expression as well (**Figure 4D, 4E**). We also found that knockdown of WNK1 decreased SMURF2 protein expression (**Figure 4F**). As expected, knockdown of SMURF2 increased ALK5 protein expression (**Figure 4G, 4H**). Interestingly, SMURF2 knockdown also resulted in a corresponding increase in the phosphorylated active form of OSR1, suggestive of enhanced WNK1 activity (**Figure 4G, 4I**). Given the regulation of ALK5 by WNK1, and that of WNK1 by SMURF2, these data suggest that the increase in ALK5 protein expression by SMURF2 knockdown may, in part, result from increased WNK1 activity. Overall, these results suggest that WNK1 kinase activity and SMURF1/2 reciprocally regulate each other to affect responses to TGF-β (**Figure 4J**).

**Figure 4:**
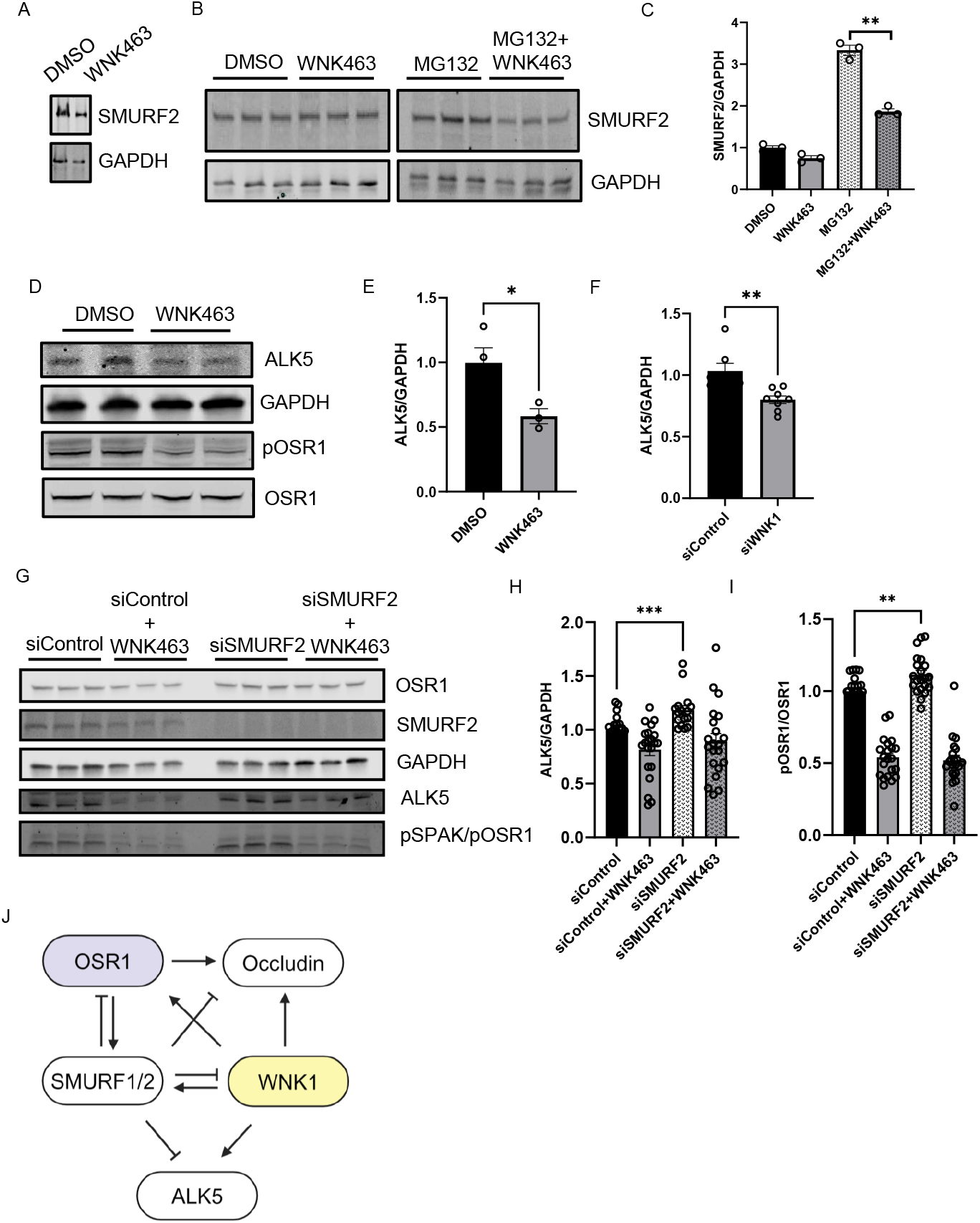
WNK1 kinase activity mediates SMURF2-dependent regulation of ALK5: **A**) Representative Western Blot showing SMURF2 protein levels upon WNK463 overnight treatment (1 µM). **B**) Representative Western Blot showing SMURF2 protein levels upon overnight WNK463 (1 µM) ± MG132 (10 µM). **C**) Corresponding quantification of ‘B’ showing decreased SMURF2 levels upon co-treatment with WNK463 and MG132; n=3. **D**) Representative Western Blot showing ALK5 levels upon DMSO or WNK463 (1 µM) in HDMECs. **E**) Corresponding quantification of ‘D’ showing decreased ALK5 levels upon WNK463 (1 µM) treatment; n=3. **F**) Quantification of ALK5 in HDMECs treated with siControl or siWNK1; n=6. **G**) Representative Western Blot showing ALK5 and pOSR1 levels upon siControl or siSMURF2 co-treated with DMSO or WNK463 (1 µM) in HDMECs. **H**) Corresponding quantification of ‘G’ showing increased ALK5 levels upon siSMURF2 treatment; n=21. **I**) Corresponding quantification of ‘G’ showing increased pOSR1 levels upon siSMURF2 treatment; n=21. **J**) Model representing inter-regulation among WNK1, SMURF1/2 and ALK5. Data are represented as Mean±SE; analyzed by unpaired two-tailed Student’s t-test or one-way ANOVA. *p<0.05, **p<0.005, ***p<0.0005.

## Discussion

Unanticipated findings revealed that WNK1 and SMURFs reciprocally regulate each other to fine tune TGF-β signaling. Aggregated structures containing SMURF1/2 and WNK1 in our study are similar to those observed with respect to SMURF2 and clustering of SMURF2 is suggested to regulate its E3 ubiquitin ligase activity. [18,26]. We show that depletion or inhibition of WNK1 decreases SMURF2. The amount of SMURF2 protein present upon inhibition of SMURF1 was dependent on WNK1 protein amount. One possible explanation is that SMURF1 regulates SMURF2 protein, and this is dependent on the relative expression of WNK1. SMURF2 inhibits its own ubiquitinase activity and is thereby stabilized. Therefore, it is possible that inhibition of WNK1 may enhance the activity of SMURF1/2 and thereby lead to downregulation of itself, and ALK5 as observed in this study. Future studies will focus on addressing the mechanistic details of this potential mode of regulation.

Our study reports that WNK1 colocalizes with SMURF1/2 in large punctate structures. SMURF1 mediates p62 biomolecular condensation to promote autophagy of NRF2, a protein that is known to interact with WNK1 to enhance cellular oxidative response [37, 38]. However, we did not find co-localization of p62 in these WNK1-SMURF1/2 large punctate structures (data not shown). The WW-domains of SMURF/NEDD4 E3 ligases generally recognize and bind proline-rich sequences such as PY-motifs on substrate proteins [26]. Interestingly, PY-motifs are also found in WNK1. Activity of E3 ligases such as SMURFs are regulated by clustering of the PY motifs on their substrates [26]. WNK1 has been reported to form punctate structures referred to as WNK bodies which are membraneless cytosolic signaling foci that sequester WNKs. These structures have been reported in kidney epithelial cells during changes in total body potassium balance [20]. Therefore, we speculate that these large punctate structures we observed in our study may be similar to the WNK bodies and that such clustering of WNK1 may regulate the activity of SMURFs to affect TGF-β signaling output. Nevertheless, this interesting hypothesis requires further examination.

We found that SMURF2 interacts with OSR1 via its conserved C-terminal (CCT) domain. The CCT domain of OSR1 binds substrates and other proteins via a short conserved RFxV motifs [34,35]. We had previously reported that OSR1 CCT interacts with RFxV motif containing proteins such as occludin [10]. We found that SMURF2 interacts with occludin and this interaction was enhanced upon inhibition with the pan-WNK inhibitor WNK463 [10], but decreased upon co-incubation with the CCT blocking peptide. These observations suggest that WNK activity regulates the interaction between WNK1/OSR1, occludin and SMURF2. We propose that CCT OSR1 mediates the interaction between SMURF2 and occludin. SMURF2 has been shown to be in a complex with occludin during TGF-β signaling [13] but the mechanistic significance of interaction between SMURF2 and occludin remains to be understood.

Signal transduction pathways regulated by TGF-β control a diverse array of cellular processes such as vascular remodeling [39]. Depending on the cellular context, TGFβ signaling pathway exhibits differential and opposing responses [39-42]. This occurs via complex regulatory mechanisms and crosstalk among several TGF-β mediators [39-42]. The duration and amplitude of TGF-β signaling are tightly controlled by various proteasome-mediated degradation mediators involving E3 ubiquitin ligases such as SMURF1/2, among others [39-42]. These observations suggest that the dynamic convergence of inputs from WNK1 to SMURFs affects TGF-β receptors, essential SMAD transcription factors and tight junction components to fine-tune TGF-β signaling to facilitate diverse cellular outcomes. Therefore, we propose that expression and activity of WNK1 contributes to the context-specificity in TGF-β signaling.

## Methods

### Cell lines

Human Umbilical Vein Endothelial Cells (HUVEC: ATCC, PCS-100-013) were grown in complete VascuLife® EnGS media kit (Fisher Scientific, 50-311-891) supplemented as per manufacturer’s instructions. In experiments requiring knockdowns, Human Dermal Microvascular Endothelial Cells (HMEC-1: ATCC, CRL-3243) were used. These cells were grown in complete MCDB media (Fisher Scientific, MT15100CV) supplemented with 10% fetal bovine serum (Sigma-Aldrich, F0926), 1% L-glutamine, 1% penicillin and streptomycin, 1μg/mL hydrocortisone (Sigma Aldrich, H0888 or H6909), and 10 ng/mL epidermal growth factor (EGF: Cell Signaling Technology, 8916SC). All cells were maintained at 37°C and 5% CO_2_.

### Co-immunoprecipitation

Cells were lysed in 1X lysis buffer (50mM HEPES, 0.1M NaCl, 0.5mM EDTA, 0.1% SDS) supplemented with protease inhibitor cocktail, PMSF, and phosphatase inhibitors (PhosStop). Cell extracts were harvested and cleared by centrifugation. 1X IP buffer (50mM HEPES, 0.1M NaCl, 0.5mM EDTA, and 1% CHAPS (Sigma Aldrich, C3023) supplemented with protease inhibitor cocktail, PMSF, and phosphatase inhibitors (Sigma Aldrich, 4906837001) was added in a 2:1 ratio to the cell lysate. Samples were incubated with primary antibody (control sample incubated with rabbit IgG primary antibody: 1:100) overnight at 4°C and then with Protein A/G PLUS-Agarose (Santa Cruz Biotechnology, sc-2003) beads for 1 hour with head-to-tail rotation. This was performed either in the absence or presence of CCT blocking peptide peptide SAGRRFIVSPVPE (100 µM) (United Biosystems). Samples were then washed three times with 1X IP buffer before adding 5X SDS buffer (0.25% bromophenol blue, 0.5M DTT, 50% glycerol, 10% SDS, 0.25M Tris-Cl) and heating at 90°C for 2 minutes. Samples were then run on 4-20% Mini-PROTEAN® TGX™ Precast Protein Gels (Bio-Rad, 4568096) or 12% polyacrylamide home-made gels before being transferred to PVDF membranes. Membranes were then washed in TBS-T before being blocked with TBS-based blocking buffer (LI-COR). Membranes were incubated with primary antibodies and then washed again before being incubated with species-specific, light chain-specific secondary antibodies (Jackson ImmunoResearch Labs, 115-655-174 and 211-622-171) and imaged using LI-COR imaging.

### Immunofluorescence

HUVEC cells were fixed on glass coverslips (Fisher Scientific, 12-545-80) with 4% paraformaldehyde for 20 minutes at room temperature. Coverslips were washed with sterile PBS and blocked in 5% normal goat serum (Life Technologies, 50-062Z) before incubating with primary antibodies for 1 hour at room temperature. Coverslips were washed with 1X PBS. Subsequently, cells were incubated with an Alexa Fluor® 488 conjugated goat-anti-mouse secondary antibody (Thermo Fisher Scientific, A11029) and Alexa Fluor® 594 conjugated goat-anti-rabbit secondary antibody (Invitrogen, A11037) for 30 min at room temperature in dark, and the slides were mounted with DAPI Fluoromount-G (Thermo Fisher Scientific, 00-4959-52). Immunofluorescence images were acquired using a Zeiss LSM880 inverted confocal microscope (Carl Zeiss, Oberkochen, Germany). Images were deconvolved using AutoQuant® software (Media Cybernetics, USA).

### siRNA knockdown

Oligonucleotides encoding siRNA for human WNK1 (siWNK1: 5’ CAGACAGUGCAGUAUUCACTT 3’), control siRNA (Thermo Fisher Scientific, 4390844) as in [99], OSR1 siRNA (Thermo Fisher Scientific, s19303 Silencer® Select), and SMURF2 siRNA (sc-41675, Santa Cruz Biotechnology). HDMEC cells were transfected with 20 nM siRNA using Lipofectamine RNAiMax reagent (Thermo Fisher Scientific, 13778150). After 24-72 hours of transfection, cells were provided with their respective treatments and were then harvested in 1X SDS buffer (0.05% bromophenol blue, 0.1M DTT, 10% glycerol, 2% SDS, and 0.05M Tris-Cl) with 5% β-mercaptoethanol.

### Immunoblotting

Cell lysates containing 1X SDS buffer were homogenized with 27-G syringe and whole lysates were run on 4-20% Mini-PROTEAN® TGX™ Precast Protein Gels (Bio-Rad, 4568096) or 6/10/12% home-made polyacrylamide gels before being transferred to PVDF membranes (Bio-Rad, 1620177). Membranes were then washed in TBS-T before being blocked with TBS-based blocking buffer (LI-COR). Membranes were incubated with primary antibodies and then washed again before being incubated with species-specific secondary antibodies and imaged and quantified using LI-COR.

### Reagents

WNK463 (Selleck Chemicals, S8358), SMURF1 inhibitor A01 (Sigma Aldrich, SML1404), MG132 (Sigma Aldrich, M7449), TGF-β1 (Cell Signaling Technology, 8915LC), anti-Vinculin antibody (Sigma Aldrich, V9131), anti-pOSR1 antibody (EMD Millipore, 07-2273), anti-OSR1 polyclonal antibody (Cell Signaling, 3729S), anti-OSR1 monoclonal antibody (VWR, 10624-616), anti-WNK1 antibody (Cell Signaling, 4979S), anti-GAPDH antibody (Cell signaling Technology, 97166L), anti-SMURF1 antibody (Santa Cruz Biotechnology, sc-100616), anti-SMURF2 antibody (Santa Cruz Biotechnology, sc-393848), anti-flag antibody (Sigma-Aldrich, F1804), Q256 WNK1 antibody was homemade as in [7], Optimem (Invitrogen, 51985-034), Lipofectamine 2000 (Life Technologies, 11668019), bumetanide (Sigma Aldrich, B3023), 96-well plates (Corning, 3904 or Greiner, 655090).

### Statistics and Reproducibility

The data are presented mean±SEM from at least three independent experiments with similar results. All presented micrographs (immunofluorescence images) are representative images from three representative experiments as indicated in the figure legends. For the quantification of immunofluorescence images, the number of cells used for each representative experiment is indicated and p values between two groups were determined using unpaired t-tests. Results are expressed as mean ± SEM. Single intergroup comparisons between 2 groups were performed with 2-tailed Student’s *t*-test as specifically mentioned in each case. p < 0.05 was considered statistically significant.

## Data sharing

We will follow all NIH policies with respect to sharing reagents, materials, and information with other investigators. Detailed protocols are provided to everyone who requests them. Upon publication, this manuscript will be submitted to the National Library of Medicine’s PubMed Central as outlined by NIH policy.

## Acknowledgements

The authors thank Joseph P. Albanesi, UT Southwestern Department of Pharmacology and members of Cobb lab for valuable suggestions, and Dionne Ware for administrative assistance. These studies were supported by NIH R01 HL147661 and Mary Kay Foundation grant 18-18 to MHC, American Heart Association postdoctoral fellowship 18POST34030438 to ABJ, CPRIT training grant RP160157 for early support of ABJ, and Welch Foundation grant I1243 to MHC. The authors would like to acknowledge the assistance of the UT Southwestern Live Cell Imaging Facility, a Shared Resource of the Harold C. Simmons Cancer Center, supported in part by an NCI Cancer Center Support Grant, 1P30 CA142543-01 and NIH Shared Instrumentation Award 1S10 OD021684-01 to Dr. Kate Luby-Phelps (LSM880 Airyscan).

## Author contribution

**ABJ**: Conceptualized, supervised, designed and performed experiments, performed analysis, wrote manuscript, generated initial figures; **DB**: performed experiments; **CW**: Analyzed data; **KLP**: Discussed data; **MHC**: Supervision, acquired funding, edited manuscript.

## References

1. David CJ, Massagué J. Contextual determinants of TGFβ action in development, immunity and cancer. Nat Rev Mol Cell Biol. 2018 Jul;19(7):419–435.

2. Xie J, Wu T, Xu K, Huang IK, Cleaver O, Huang CL. Endothelial-specific expression of WNK1 kinase is essential for angiogenesis and heart development in mice. Am J Pathol. 2009 Sep;175(3):1315–27.

3. Xie J, Yoon J, Yang SS, Lin SH, Huang CL. WNK1 protein kinase regulates embryonic cardiovascular development through the OSR1 signaling cascade. J Biol Chem. 2013 Mar 22;288(12):8566–8574.

4. Xu B, English JM, Wilsbacher JL, Stippec S, Goldsmith EJ, Cobb MH. WNK1, a novel mammalian serine/threonine protein kinase lacking the catalytic lysine in subdomain II. J Biol Chem. 2000 Jun 2;275(22):16795–801.

5. Vitari AC, Deak M, Morrice NA, Alessi DR. The WNK1 and WNK4 protein kinases that are mutated in Gordon’s hypertension syndrome phosphorylate and activate SPAK and OSR1 protein kinases. Biochem J. 2005 Oct 1;391(Pt 1):17–24.

6. Moriguchi T, Urushiyama S, Hisamoto N, Iemura S, Uchida S, Natsume T, Matsumoto K, Shibuya H. WNK1 regulates phosphorylation of cation-chloride-coupled cotransporters via the STE20-related kinases, SPAK and OSR1. J Biol Chem. 2005 Dec 30;280(52):42685–93.

7. Xu BE, Stippec S, Chu PY, Lazrak A, Li XJ, Lee BH, English JM, Ortega B, Huang CL, Cobb MH. WNK1 activates SGK1 to regulate the epithelial sodium channel. Proc Natl Acad Sci U S A. 2005 Jul 19;102(29):10315–20.

8. Anselmo AN, Earnest S, Chen W, Juang YC, Kim SC, Zhao Y, Cobb MH. WNK1 and OSR1 regulate the Na+, K+, 2Cl-cotransporter in HeLa cells. Proc Natl Acad Sci U S A. 2006 Jul 18;103(29):10883–8.

9. Taylor CA 4th, An SW, Kankanamalage SG, Stippec S, Earnest S, Trivedi AT, Yang JZ, Mirzaei H, Huang CL, Cobb MH. OSR1 regulates a subset of inward rectifier potassium channels via a binding motif variant. Proc Natl Acad Sci U S A. 2018 Apr 10;115(15):3840–3845.

10. Jaykumar AB, Plumber S, Barry DM, Binns D, Wichaidit C, Grzemska M, Earnest S, Goldsmith EJ, Cleaver O, Cobb MH. WNK1 collaborates with TGF-β in endothelial cell junction turnover and angiogenesis. Proc Natl Acad Sci U S A. 2022 Jul 26;119(30):e2203743119.

11. Lee BH, Chen W, Stippec S, Cobb MH. Biological cross-talk between WNK1 and the transforming growth factor beta-Smad signaling pathway. J Biol Chem. 2007 Jun 22;282(25):17985–17996.

12. Ozdamar B, Bose R, Barrios-Rodiles M, Wang HR, Zhang Y, Wrana JL. Regulation of the polarity protein Par6 by TGFbeta receptors controls epithelial cell plasticity. Science. 2005 Mar 11;307(5715):1603–9.

13. Barrios-Rodiles M, Brown KR, Ozdamar B, Bose R, Liu Z, Donovan RS, Shinjo F, Liu Y, Dembowy J, Taylor IW, Luga V, Przulj N, Robinson M, Suzuki H, Hayashizaki Y, Jurisica I, Wrana JL. High-throughput mapping of a dynamic signaling network in mammalian cells. Science. 2005 Mar 11;307(5715):1621–5.

14. Murakami T, Felinski EA, Antonetti DA. Occludin phosphorylation and ubiquitination regulate tight junction trafficking and vascular endothelial growth factor-induced permeability. J Biol Chem. 2009 Jul 31;284(31):21036–46.

15. Wong V. Phosphorylation of occludin correlates with occludin localization and function at the tight junction. Am J Physiol. 1997 Dec;273(6):C1859–67.

16. Du D, Xu F, Yu L, Zhang C, Lu X, Yuan H, Huang Q, Zhang F, Bao H, Jia L, Wu X, Zhu X, Zhang X, Zhang Z, Chen Z. The tight junction protein, occludin, regulates the directional migration of epithelial cells. Dev Cell. 2010 Jan 19;18(1):52–63

17. Liu X, Dreffs A, Díaz-Coránguez M, Runkle EA, Gardner TW, Chiodo VA, Hauswirth WW, Antonetti DA. Occludin S490 Phosphorylation Regulates Vascular Endothelial Growth Factor-Induced Retinal Neovascularization. Am J Pathol. 2016 Sep;186(9):2486–99.

18. Kavsak P, Rasmussen RK, Causing CG, Bonni S, Zhu H, Thomsen GH, Wrana JL. Smad7 binds to Smurf2 to form an E3 ubiquitin ligase that targets the TGF beta receptor for degradation. Mol Cell. 2000 Dec;6(6):1365–75.

19. Ebisawa T, Fukuchi M, Murakami G, Chiba T, Tanaka K, Imamura T, Miyazono K. Smurf1 interacts with transforming growth factor-beta type I receptor through Smad7 and induces receptor degradation. J Biol Chem. 2001 Apr 20;276(16):12477–80.

20. Boyd-Shiwarski CR, Shiwarski DJ, Roy A, Namboodiri HN, Nkashama LJ, Xie J, McClain KL, Marciszyn A, Kleyman TR, Tan RJ, Stolz DB, Puthenveedu MA, Huang CL, Subramanya AR. Potassium-regulated distal tubule WNK bodies are kidney-specific WNK1 dependent. Mol Biol Cell. 2018 Feb 15;29(4):499–509.

21. Koefoed K, Skat-Rørdam J, Andersen P, Warzecha CB, Pye M, Andersen TA, Ajbro KD, Bendsen E, Narimatsu M, Vilhardt F, Pedersen LB, Wrana JL, Anderson RH, Møllgård K, Christensen ST, Larsen LA. The E3 ubiquitin ligase SMURF1 regulates cell-fate specification and outflow tract septation during mammalian heart development. Sci Rep. 2018 Jun 22;8(1):9542

22. Lin X, Liang M, Feng XH. Smurf2 is a ubiquitin E3 ligase mediating proteasome-dependent degradation of Smad2 in transforming growth factor-beta signaling. J Biol Chem. (2000) 275:36818–22.

23. Koganti P, Levy-Cohen G, Blank M. Smurfs in Protein Homeostasis, Signaling, and Cancer. Front Oncol. 2018 Aug 2;8:295.

24. Zhang Y, Chang C, Gehling DJ, Hemmati-Brivanlou A, Derynck R. Regulation of Smad degradation and activity by Smurf2, an E3 ubiquitin ligase. Proc Natl Acad Sci U S A. 2001 Jan 30;98(3):974–9.

25. Tang LY, Yamashita M, Coussens NP, Tang Y, Wang X, Li C, Deng CX, Cheng SY, Zhang YE. Ablation of Smurf2 reveals an inhibition in TGF-β signalling through multiple monoubiquitination of Smad3. EMBO J. 2011 Nov 1;30(23):4777–89.

26. Mund T, Pelham HR. Substrate clustering potently regulates the activity of WW-HECT domain-containing ubiquitin ligases. J Biol Chem. 2018 Apr 6;293(14):5200–5209.

27. Wang HR, Ogunjimi AA, Zhang Y, Ozdamar B, Bose R, Wrana JL. Degradation of RhoA by Smurf1 ubiquitin ligase. Methods Enzymol. 2006;406:437–47.

28. McCormick JA, Yang CL, Zhang C, Davidge B, Blankenstein KI, Terker AS, Yarbrough B, Meermeier NP, Park HJ, McCully B, West M, Borschewski A, Himmerkus N, Bleich M, Bachmann S, Mutig K, Argaiz ER, Gamba G, Singer JD, Ellison DH. Hyperkalemic hypertension-associated cullin 3 promotes WNK signaling by degrading KLHL3. J Clin Invest. 2014 Nov;124(11):4723–36.

29. Jung JU, B Ghosh A, Earnest S, Deaton SL, Cobb MH. UBR5 is a novel regulator of WNK1 stability. Am J Physiol Cell Physiol. 2022 Jun 1;322(6):C1176–C1186.

30. Roy A, Al-Qusairi L, Donnelly BF, Ronzaud C, Marciszyn AL, Gong F, Chang YP, Butterworth MB, Pastor-Soler NM, Hallows KR, Staub O, Subramanya AR. Alternatively spliced proline-rich cassettes link WNK1 to aldosterone action. J Clin Invest. 2015 Sep;125(9):3433–48.

31. Al-Qusairi L, Basquin D, Roy A, Stifanelli M, Rajaram RD, Debonneville A, Nita I, Maillard M, Loffing J, Subramanya AR, Staub O. Renal tubular SGK1 deficiency causes impaired K+ excretion via loss of regulation of NEDD4-2/WNK1 and ENaC. Am J Physiol Renal Physiol. 2016 Aug 1;311(2):F330–42.

32. Roy A, Al-Qusairi L, Donnelly BF, Ronzaud C, Marciszyn AL, Gong F, Chang YP, Butterworth MB, Pastor-Soler NM, Hallows KR, Staub O, Subramanya AR. Alternatively spliced proline-rich cassettes link WNK1 to aldosterone action. J Clin Invest. 2015 Sep;125(9):3433–48.

33. Kuratomi G, Komuro A, Goto K, Shinozaki M, Miyazawa K, Miyazono K, Imamura T. NEDD4-2 (neural precursor cell expressed, developmentally down-regulated 4-2) negatively regulates TGF-beta (transforming growth factor-beta) signalling by inducing ubiquitin-mediated degradation of Smad2 and TGF-beta type I receptor. Biochem J. 2005 Mar 15;386(Pt 3):461–70.

34. Delpire E, Gagnon KB. Genome-wide analysis of SPAK/OSR1 binding motifs. Physiol Genomics. 2007 Jan 17;28(2):223–31.

35. Taylor CA IV, Jung J, Kankanamalage SG, Li J, Grzemska M, Jaykumar AB, Earnest S, Stippec S, Saha P, Sauceda E, Cobb MH. Predictive and Experimental Motif Interaction Analysis Identifies Functions of the WNK-OSR1/SPAK Pathway. bioRxiv 2024;06.26.600905.

36. Cao Y, Wang C, Zhang X, Xing G, Lu K, Gu Y, He F, Zhang L. Selective small molecule compounds increase BMP-2 responsiveness by inhibiting Smurf1-mediated Smad1/5 degradation. Sci Rep. 2014 May 14;4:4965.

37. Li L, Xie D, Yu S, Ma M, Fan K, Chen J, Xiu M, Xie K, Li Y, Yong G. WNK1 Interaction with KEAP1 Promotes NRF2 Stabilization to Enhance the Oxidative Stress Response in Hepatocellular Carcinoma. Cancer Res. 2024 Jun 17.

38. Xia Q, Li Y, Xu W, Wu C, Zheng H, Liu L, Dong L. Enhanced liquidity of p62 droplets mediated by Smurf1 links Nrf2 activation and autophagy. Cell Biosci. 2023 Feb 21;13(1):37

39. Lebrin F, Deckers M, Bertolino P, Ten Dijke P. TGF-beta receptor function in the endothelium. Cardiovasc Res. 2005 Feb 15;65(3):599–608.

40. Bellomo C, Caja L, Moustakas A. Transforming growth factor β as regulator of cancer stemness and metastasis. Br J Cancer. 2016 Sep 27;115(7):761–9.

41. Nakagawa T, Li JH, Garcia G, Mu W, Piek E, Böttinger EP, Chen Y, Zhu HJ, Kang DH, Schreiner GF, Lan HY, Johnson RJ. TGF-beta induces proangiogenic and antiangiogenic factors via parallel but distinct Smad pathways. Kidney Int. 2004 Aug;66(2):605–13.

42. Attisano L, Wrana JL. Signal transduction by the TGF-beta superfamily. Science. 2002 May 31;296(5573):1646–7.

